# Self-Organizing Assembloids Reveal Enteric Nervous System Dynamics in Gut Homeostasis and Regeneration

**DOI:** 10.1101/2025.01.14.632538

**Authors:** Ilke Sari, Beliz Uzun, Melda O. Oguz, Eren Gursoy, Nurmuhammet Satlykov, Dila A. Gemalmayan, Nagihan Gonullu, Tomas Valenta, Fatima Aerts Kaya, Merve Gizer, Petek Korkusuz, Bahar Degirmenci

## Abstract

The enteric nervous system (ENS) is essential for intestinal health, exhibiting adaptability to environmental and physiological challenges. However, the mechanisms underlying ENS plasticity and resilience remain poorly understood. Organoid technology has revolutionized in vitro modeling by accurately replicating epithelial structures and enabling significant advancements in understanding gastrointestinal biology. However, traditional organoids are limited in their ability to study the ENS, as they lack the multicellular composition and functional architecture necessary to model complex interactions between neurons, glia, mesenchymal, smooth muscle, and epithelial cells. To address these limitations, we developed murine ENS-Rich Assembloids (ERAs) that self-organize to replicate the cellular diversity, including the epithelial structure, and functional architecture of native colonic tissue. These assembloids recreate neuron-glia interactions, reflect regenerative processes, and provide a novel platform for studying ENS dynamics under controlled conditions. Integrating findings from assembloids and an in vivo murine model, we demonstrate that inflammation induces coordinated reorganization of S100b+ glial cells, TUJ1+ neurons, PDGFRA+ mesenchymal cells, and epithelial cells, revealing conserved mechanisms of ENS plasticity. We identify pleiotrophin (PTN) signaling via Protein Tyrosine Phosphatase Receptor Type Z1 (PTPRZ1) as a key pathway facilitating neural elongation and enhancing neuron-glia interactions. Moreover, we show that activated neurons transfer lipids to glial cells, revealing a novel support mechanism during inflammation. These findings position enteric glia as protective hubs for neurons, fostering ENS adaptability and tissue regeneration. By building on the foundational success of organoid technology and addressing its limitations for studying the ENS, ENS-rich assembloids establish a transformative tool for investigating ENS responses in health, disease, and tissue repair.

## Introduction

The enteric nervous system exhibits remarkable capacity for regeneration and plasticity, adapting to injury, inflammation, and physiological stress^1-4^. This plasticity is driven by dynamic interactions between enteric neurons and glial cells, which play integral roles in neuroprotection, tissue homeostasis, and repair^5-13^. Enteric glial cells, in particular, act as modulators of neuroactive substances, regulators of inflammatory responses, and contributors to adult neurogenesis^14-22^. Despite their importance, the mechanisms by which glial cells support neural survival and ENS function remain elusive. Unraveling these processes is essential for advancing gastrointestinal health and developing therapies for inflammatory and degenerative conditions. Current in vitro models, such as adult stem cell derived organoids, fail to replicate the complexity of native ENS interactions^23-26^. Efforts to overcome these constraints include coculturing pluripotent stem cell derived organoids with neural crest cells, leading to the formation of pluripotent stem cell derived intestinal tissue featuring a functional ENS^27^. However, the in vivo reliance for ENS formation diminishes the advantages of these systems as controllable in vitro platforms. Recent advancements, such as the generation of gastrointestinal assembloids, have illuminated the interplay between epithelial crypts and their mesenchymal niche^28^. These studies underscore the critical role of mesenchymal cells in guiding epithelial organization and emphasize the importance of multicellular interactions in tissue dynamics. Despite these advances, fully integrating an ENS compartment into such systems remains a significant challenge.

To address this limitation, we developed murine ENS-rich assembloids (ERAs)—a three-dimensional in vitro model incorporating ENS cells, mesenchymal cells, smooth muscle cells (SMCs) and epithelial cells. The ERAs self-organize into structures that closely mimic native colonic tissue, offering a robust platform to study ENS behavior under physiological conditions. Using the ERAs alongside an in vivo model, we explored ENS plasticity during inflammation and recovery. Furthermore, we identified the PTN-PTPRZ1 signaling axis as a key regulator of neural elongation and neuron-glia interactions. We also uncover a novel mechanism in which activated neurons transfer lipids to glial cells, acting as critical protective nodes for neurons. These insights establish ERAs as a transformative tool for studying ENS plasticity and position them at the forefront of regenerative and therapeutic research.

## Results

### Development of Enteric Nervous System-Rich Assembloids

Organoids derived from adult stem cells have revolutionized in vitro modeling by providing platforms to investigate mammalian physiology, development, and disease^29^. However, these models predominantly replicate epithelial structures and fail to capture the multicellular complexity of native tissues, including the ENS and SMC components, which are critical for organ function. To overcome this limitation, we developed a more complex model, ERA. In constructing this model, we have selected specific cell types for their complementary roles: mesenchymal cells provide structural support and guide epithelial organization^30-36^, ENS cells regulate cellular alignment and communication^37,38^, and SMCs contribute to contractility and tissue integrity^39^.

Remarkably, the ERAs self-organized into colonic tissue structures without external guidance, reflecting the intrinsic ability of these cells to mimic native tissue architecture and interactions (Fig 1a, Extended Data Fig. 1a). Histological analysis confirmed the presence of key cell populations. E-cadherin marked epithelial cells forming crypt-like structures, while mesenchymal cells (PDGFRA-positive) were distributed along the crypts and enveloped the epithelial layer, replicating the organization of native colonic tissue (Fig. 1b, Extended Data Fig. 1b). Neural organization within the assembloid closely resembled that of native tissue. ENS cells, identified by TUJ1 and S100B, were observed aligning with the epithelial and mesenchymal layers, contributing to neural organization (Fig. 1b-d, Extended Data Fig. 1b). PHOX2B-positive cells near the crypts co-expressed HuC-D, confirming their neural identity (Fig. 1e). SMCs, marked by ACTA2, formed contractile layers that added structural and functional fidelity to the assembloid (Fig. 1f). Proliferation was evident in both epithelial and mesenchymal compartments. Ki67-positive cells were localized at the crypt bases, co-expressing E-cadherin, indicating active epithelial renewal (Fig. 1g). PDGFRA-positive mesenchymal cells in the muscle region also expressed Ki67, highlighting their dual role in structural support and tissue development (Extended Data Fig. 1c).

**Figure 1.**
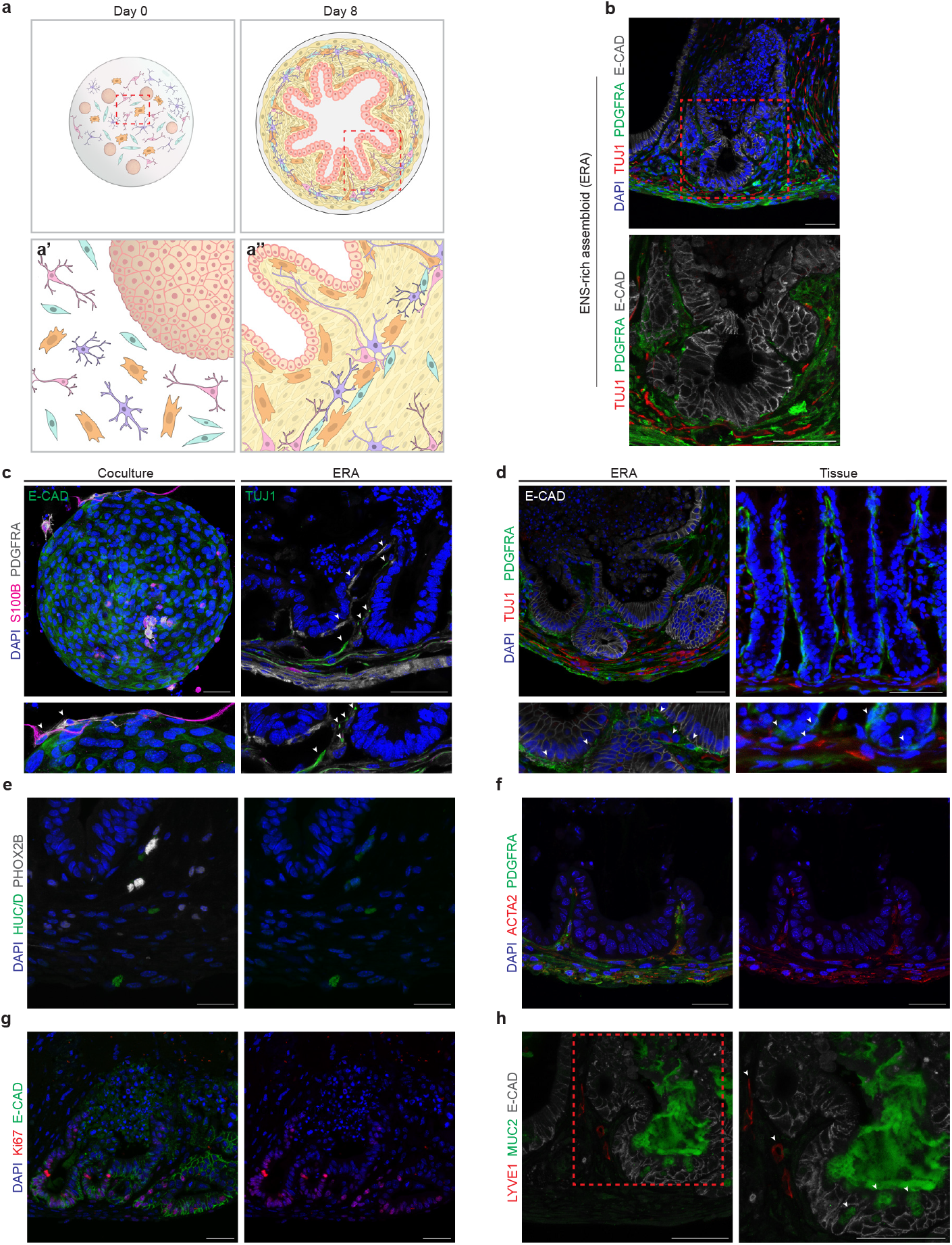
ENS-rich assembloids (ERAs) effectively mirror the cellular diversity and native physiological organization of the colon. **(a)** Illustration of ENS-rich assembloid (ERA) droplet in day 0. Inset (a’) shows that the cells were randomly mixed within the Matrigel. By Day 8, the assembloid droplet demonstrates self-organization of diverse cell types, forming crypt-like structures and tissue-like architecture including smooth muscle, enteric nervous system (ENS) and mesenchymal cells. Inset (a’’) highlights the coordinated cell movement of mesenchymal, glial, and neuronal cells along smooth muscle cells. **(b)** ERAs were analyzed via immunohistochemistry, showing tissue-like organization with epithelial marker E-Cadherin (E-CAD), enteric neuron marker TUJ1, and mesenchymal cell marker PDGFRA. PDGFRA+ mesenchymal cells closely surround the crypts, supporting crypt formation. **(c)** Comparison of conventional coculture and ENS-rich assembloid system. In the conventional coculture system, E-CAD+ organoids, S100B+ (glial), and PDGFRA+ mesenchymal cells surround the organoid, as shown in the inset. In contrast, in the ENS-rich assembloid, TUJ1+ enteric neurons position themselves through the invagination of crypts. Insets illustrate how glial, neuronal, and mesenchymal cells are positioned near the epithelial cell compartments, highlighting the coordinated self-organization of these cell types in the assembloid system. **(d)** Immunohistochemistry analysis for E-Cadherin, TUJ1, and PDGFRA in both assembloids and colon tissue reveals a similar spatial organization, indicated by arrowheads. **(e)** Different types of enteric neurons in the assembloids are observed as HUCD+/PHOX2B+ and HUCD+ cells. **(f)** Histological analysis of smooth muscle cell marker ACTA2 and PDGFRA+ cells reveal the presence of smooth muscle cells at the bottom of the crypts and within crypt extensions. **(g)** Immunohistochemistry analysis of Ki67 and E-CAD in assembloids shows that crypt bases express the proliferation marker Ki67. **(h)** The assembloid system accommodates multiple cell types, as demonstrated by immunohistochemistry for goblet cells (MUC2) and endothelial cells (LYVE1). Goblet cells (MUC2) secrete mucus into the assembloid lumen, while endothelial cells are localized near the crypts, forming tissue-like structures. At least three biological replicates (n=3) were used for tissue and assembloid staining (three assembloids per experimental setup). Scale bar: 50 µm.

These findings demonstrate coordinated interactions between cell types, with mesenchymal and ENS cells playing critical roles in guiding epithelial and smooth muscle organization. Further characterization revealed the presence of additional functional cell types, including goblet cells (MUC2-positive), responsible for mucus secretion, and endothelial cells (LYVE1-positive), positioned near the epithelium with morphologies consistent with native tissue (Fig. 1h). This diversity underscores the ability of the assembloid to self-organize into distinct, physiologically relevant compartments that mirror the cellular hierarchy of the native colon. The self-organizing behavior of the assembloid highlights the intrinsic capacity of these cells to form a highly structured and functional environment without external guidance. Each cell type autonomously guided its positioning and contributed to the assembly of a model that recapitulates the architecture and dynamics of native tissue. This intrinsic organization may parallel tissue repair and early developmental processes, where cell-autonomous signaling and spatial cues govern tissue assembly.

### DSS-Induced Damage Impacts Colonic Crypt Integrity and Activates Enteric Nervous System

ERAs demonstrated the intrinsic self-organizing behavior of colonic tissues, with mesenchymal, ENS, and epithelial cells autonomously forming physiologically relevant compartments. To determine whether the intrinsic self-organizing behaviors observed in ERAs are conserved during in vivo tissue repair, we investigated the interplay of ENS and mesenchymal cells during colonic recovery in a dextran sodium sulfate (DSS) induced murine damage model.

DSS treatment caused significant crypt damage and a marked reduction in proliferative stem cells in the no recovery group. In contrast, the recovery group exhibited increased expression of regeneration markers, suggesting active epithelial repair. By day five, crypt integrity was severely compromised, with proliferative progenitor cells absent, highlighting the model’s utility for studying inflammation and regeneration (Extended Data Fig. 2a, b). Damage triggered an increase in S100b+ glial cells in the muscularis externa, correlating with muscle hypertrophy in the no recovery group (Fig. 2a, b). Glial cell levels normalized during recovery (Extended Data Fig. 2c). GFAP+ glial cells localized predominantly to the myenteric plexus, with some co-expressing VIMENTIN (Extended Data Fig. 2d). PDGFRA+ mesenchymal cells were distributed across colonic layers, concentrated near enteric ganglia in the submucosal and myenteric plexuses, suggesting dynamic interactions with ENS components (Fig. 2a, b).

**Figure 2.**
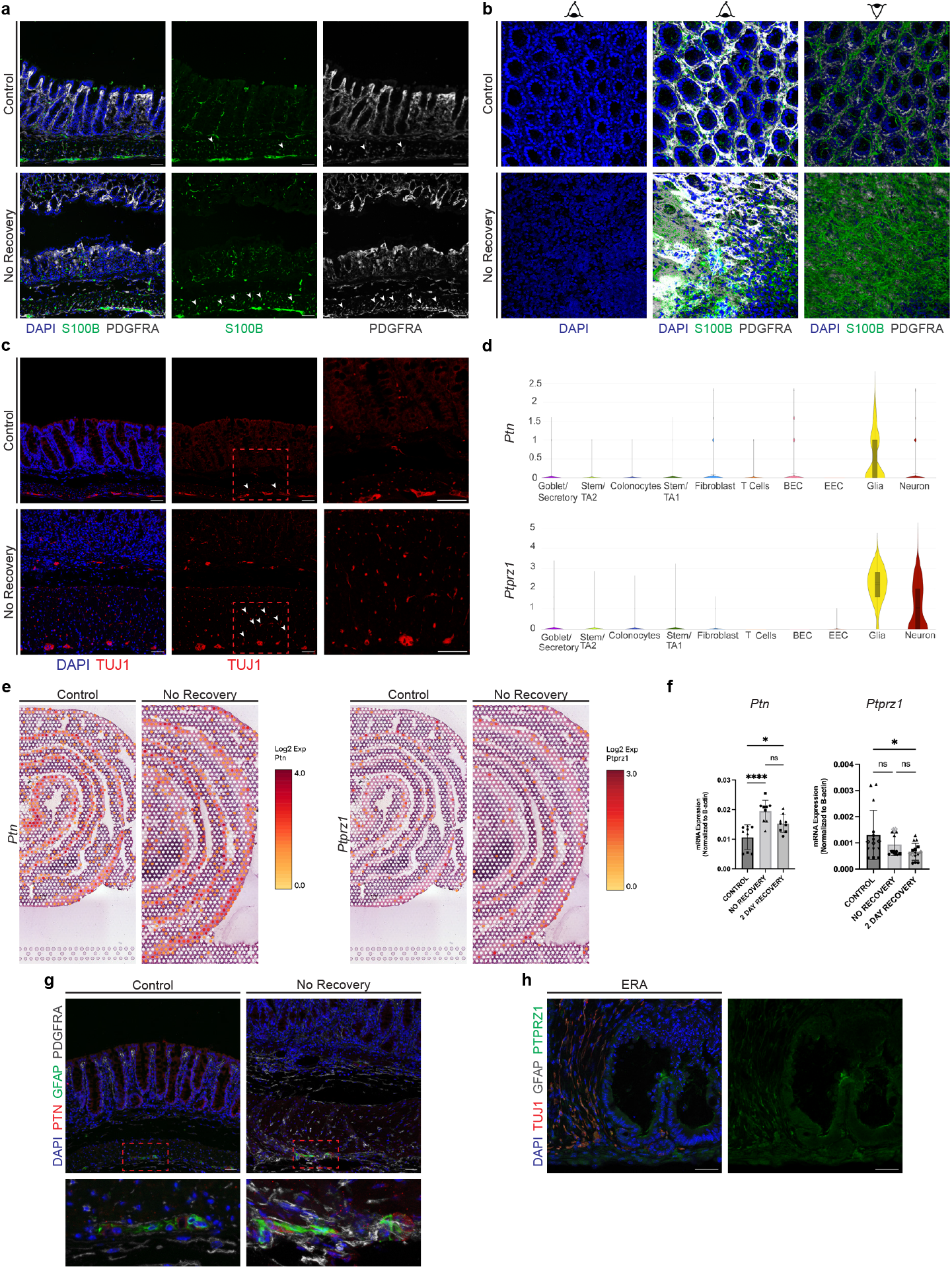
ENS cells, along with mesenchymal cells, extend towards the submucosa via the PTN-PTPRZ1 axis during damage and regeneration. **(a)** Immunohistochemistry analysis of S100B (glia) and PDGFRA (mesenchymal cells) in healthy and damaged colon. Insets show the inclination of cells from the myenteric plexus to the submucosal plexus, indicated by arrowheads. **(b)** Whole-mount immunohistochemistry of colon tissue from healthy and damaged states, showing S100B (glia) and PDGFRA (mesenchymal cell) markers. **(c)** Histological analysis of TUJ1 (neuron) in healthy and damaged colon. Insets highlight the inclination of cells from the myenteric plexus to the submucosal plexus, with arrowheads showing the direction. **(d)** Violin plots of *Ptn* and *Ptprz1* expression across different cell types, based on single-cell RNA sequencing data (Broad Institute Single Cell Portal Accession Number SCP1038). **(e)** Spatial transcriptomics analysis (GEO accession number GSE169749) of control and recovery animal models, showing *Ptn* and *Ptprz1* expression. Each spot indicates expression levels, with higher expression in muscle regions. **(f)** qRT-PCR analysis of *Ptn* and *Ptprz1* of control, no recovery, and 2-day recovery colon samples. Data are presented as mean ± SD with statistical analysis via ordinary one-way ANOVA and Tukey’s multiple comparisons (*p < 0.05, **p < 0.01, ****p < 0.0001). **(g)** Histological analysis of PTN, GFAP (glia), and PDGFRA in healthy and damaged colon. **(h)** Co-staining of PTPRZ1, GFAP (glia), and TUJ1 (neuron). At least three biological replicates (n=3) were used for qRT-PCR analysis, tissue and assembloid staining (three assembloids per experimental setup). Scale bar: 50 µm.

Spatial analysis revealed that S100b+ glial cells and PDGFRA+ mesenchymal cells reoriented toward the mucosa during inflammation, indicating a coordinated response to damage. This alignment was particularly evident in no recovery and early recovery samples (Fig. 2a, b). Enteric neurons (TUJ1-positive) extended from the myenteric plexus to the submucosa, reflecting increased elongation during inflammation (Fig. 2c). S100b+ glial cells, TUJ1+ neurons, and PDGFRA+ cells formed a structural continuum that is elevated during damage, which reverted to baseline upon recovery, demonstrating the reversible nature of these structural adaptations (Fig. 2a, 2c and Extended Data Fig. 2e-g). Notably, crypt integrity and neural elongation mirrored the regenerative dynamics observed in the assembloids. This reorganization underscores a conserved mechanism of glial guidance and mesenchymal support during tissue repair. Glial cells emerged as critical hubs, consistent with the recent identification of their role in directing epithelial and neural regeneration^40^.

### PTN-PTPRZ1 Signaling Facilitates ENS Cell Elongation During Recovery

To elucidate the mechanisms driving ENS elongation, we performed independent and detailed analyses of single-cell RNA sequencing^41^ (scRNA-seq) and spatial transcriptomics^42^. We identified *Ptn* and *Ptprz1* as pivotal regulators of ENS reorganization (Fig. 2d). PTN-PTPRZ1 signaling implied to enhance cell adhesion, survival and migration^43-45^. *Ptn* was predominantly expressed in glial cells, whereas *Ptprz1* was expressed in both glia and neurons, suggesting their complementary roles in cell migration and growth (Fig. 2d). These findings further confirmed by spatial transcriptomics which localized *Ptn* expression to the muscle layer containing the two main ENS plexuses, while *Ptprz1* was enriched in the myenteric plexus. Notably, *Ptn* expression increased significantly by day 14 of recovery, whereas *Ptprz1* levels remained stable or slightly decreased, which were further confirmed by transcriptional analysis (Fig. 2e, f). This upregulation of *Ptn* highlights its critical role in directing ENS cell elongation and migration toward damaged areas, orchestrating coordinated cellular responses during tissue repair. Co-expression analysis further revealed an increased overlap of *Ptn* and *Ptprz1* expression in the myenteric plexus during recovery, underscoring their synergistic involvement in ENS cell migration and structural reorganization (Extended Data Fig. 2h).

Histological analysis confirmed PTN localization in glial cells within the myenteric ganglia, with significant upregulation following DSS treatment (Fig. 2g). Additionally, we observed clear expression of PTPRZ1 in enteric neurons within our assembloid models (Fig. 2h), supporting its role in neural structure formation and regeneration. Beyond its role in the ENS, the PTPRZ1 axis has been implicated in tissue-specific processes across multiple systems. In the central nervous system (CNS), PTPRZ1 regulates neural differentiation and glial responses to injury, primarily through interactions with its ligand PTN^46^. These interactions modulate downstream signaling pathways to mediate cytoskeletal reorganization and cell adhesion. Similarly, PTPRZ1 has been shown to influence angiogenesis and extracellular matrix remodeling, highlighting its versatility in regulating cellular migration and tissue integrity^47^. Our data reveals that PTPRZ1 extends its functional role to the ENS, mirroring its regulatory functions in the CNS. The stability of *Ptprz1* expression during recovery, coupled with the dynamic upregulation of *Ptn*, suggests a context-dependent interaction that enables precise control over cellular behaviors required for tissue repair. Genes involved in cell adhesion and migration, such as *L1cam, Mcam, Ncam1*, and *Nrxn1*, were also upregulated during recovery, aligning with processes of ENS elongation and structural reorganization (Extended Data Fig. 3a-d). Notably, the *Pdgfa-Pdgfra* axis exhibited co-expression, underscoring its critical role in mesenchymal migration and its integration into ENS recovery pathways (Extended Data Fig. 3e).

Together, these findings position the PTN-PTPRZ1 axis as a central mechanism driving ENS elongation and tissue repair. By integrating glial and neural responses to damage, this axis directs cellular migration and reorganization during recovery. The involvement of additional adhesion and migration pathways emphasizes the complexity of ENS regeneration and reflects broader roles of PTPRZ1 across diverse tissues. However, while PTN-PTPRZ1 signaling may ensure structural reorganization, metabolic stability during these processes is equally critical for sustained repair.

### Lipid Metabolism Alterations and Neural Lipid Transfer in the ENS

A detailed analysis of scRNA-seq dataset^41^ revealed genes associated with lipid metabolism. Additionally, transcriptional analysis from muscularis externa revealed upregulation of key lipid-related genes, including *Ptgs2, ApoE, Acaa2*, and *Pgd2s*, while expression of *15-PGDH*, a critical enzyme in prostaglandin degradation, was reduced (Extended Data Fig. 4a, b). Inflammation significantly impacted lipid metabolism within the ENS. These findings suggest that lipid metabolism alterations in the ENS may serve as an adaptive mechanism to protect neural integrity and maintain functionality during inflammatory stress. Specifically, glial cells may act as a lipid reservoir, buffering metabolic demands to safeguard neurons during recovery.

Enteric neurons located in myenteric plexus showed APOE expression (Fig. 3a). Also, enteric neurons within assembloids demonstrated clear APOE positivity (Fig. 3b). Furthermore, ACOT7^49^, a pivotal enzyme in fatty acid metabolism, was prominently expressed in elongating ENS cells (Extended Data Fig. 4c, d), highlighting a potential role in supporting neural-glial metabolic interactions during tissue regeneration. These findings are in line with mechanisms observed in the central nervous system (CNS), where glial cells serve as metabolic nodes. Similar to enteric glia, astrocytes protect neurons from lipids, during periods of stress, emphasizing a conserved neuroprotective mechanism across the nervous systems^48^.

**Figure 3.**
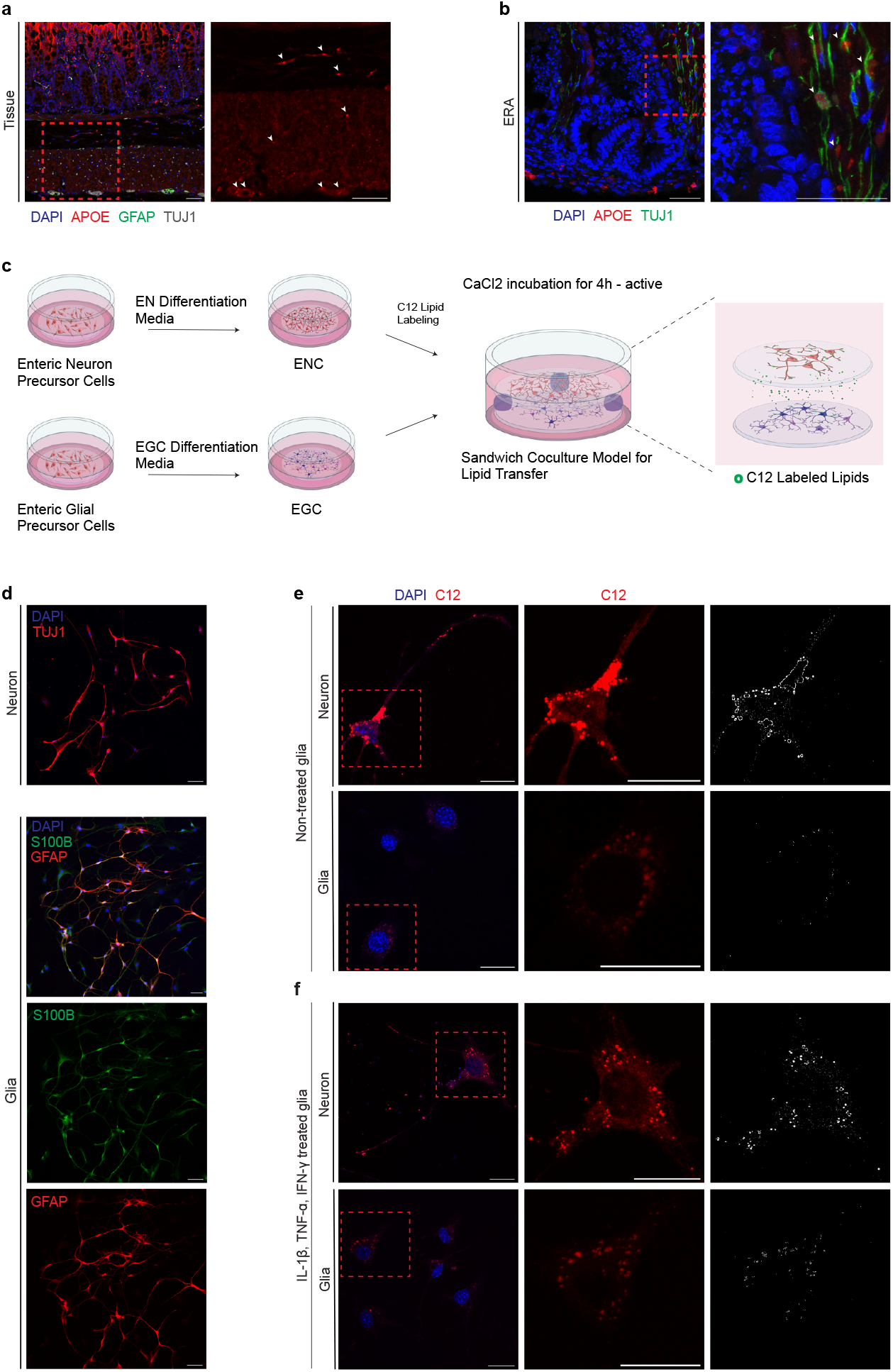
Enteric glia protect enteric neurons from lipid-induced toxicity via lipid uptake. **(a)** Histological analysis of lipid-binding protein APOE in the healthy colon, with markers for enteric neurons (TUJ1), glia (GFAP), and mesenchymal cells (PDGFRA). Inset and arrowheads highlight APOE expression in the ENS within the muscle region, submucosal, and myenteric plexus. **(b)** APOE expression in TUJ1+ enteric neurons in assembloid cultures. **(c)** Experimental scheme for lipid transfer between isolated primary enteric neurons and glia. **(d)** Histological analysis of differentiated neurons and glia using TUJ1, GFAP, and S100B (glia) markers to assess purity of cultures. **(e)** Lipid droplets (C12) observed in neurons and glia. Scale bar: 20 µm. **(f)** Lipid droplets (C12) in neurons and IL-1β, TNF-α, IFN-γ treated glia. Scale bar: 20 µm. All experiments were performed with a minimum of three biological replicates (n=3). Scale bar: 50 µm, unless indicated otherwise.

Lipid transfer assays illuminated a dynamic lipid exchange between enteric neurons and glial cells isolated from murine myenteric plexus (Fig. 3c, d). Active enteric neurons transferred fluorescently labelled lipids to glial cells, a process amplified under inflammatory conditions (Fig. 3e, f and Extended Data Fig. 5a, b). This lipid exchange appears to mitigate metabolic stress in hyperactive neurons, enabling the ENS to maintain stability during inflammatory episodes and subsequent recovery phases. Such lipid exchange is reminiscent of processes in the CNS, where neural-glial interactions facilitate lipid and metabolite transfer to maintain homeostasis^48^. This parallel highlight a shared strategy where glial cells protect neurons from metabolic overload to sustain nervous system function under stress.

These findings underscore the critical interplay between structural and metabolic adaptations in ENS remodeling. While the PTN-PTPRZ1 axis drives cellular migration and structural reorganization, lipid metabolism alterations provide essential metabolic support. Together, these pathways emphasize the complexity and coordination required for effective ENS reorganization under perturbed conditions.

### Developmental and Regenerative Parallels in ENS Organization

During early murine postnatal colon development, ENS cells exhibited coordinated migration into the lamina propria which is consistent with the previous findings in the small intestine^6^. At postnatal day 0 (P0), enteric neurons were primarily confined to the myenteric plexus, while glial cells showed limited localization beyond this region. By postnatal day 10 (P10), both enteric neurons and glial cells had extended into the submucosa, aligning within the lamina propria (Fig. 4a). At P10, HuC/D+ neurons began appearing in the submucosal region (Fig. 4b). Notably, PTN expression was consistently detected in cells within the myenteric plexus at both stages (Fig. 4b). This pattern suggests a temporal regulation of neural differentiation and migration.

**Figure 4.**
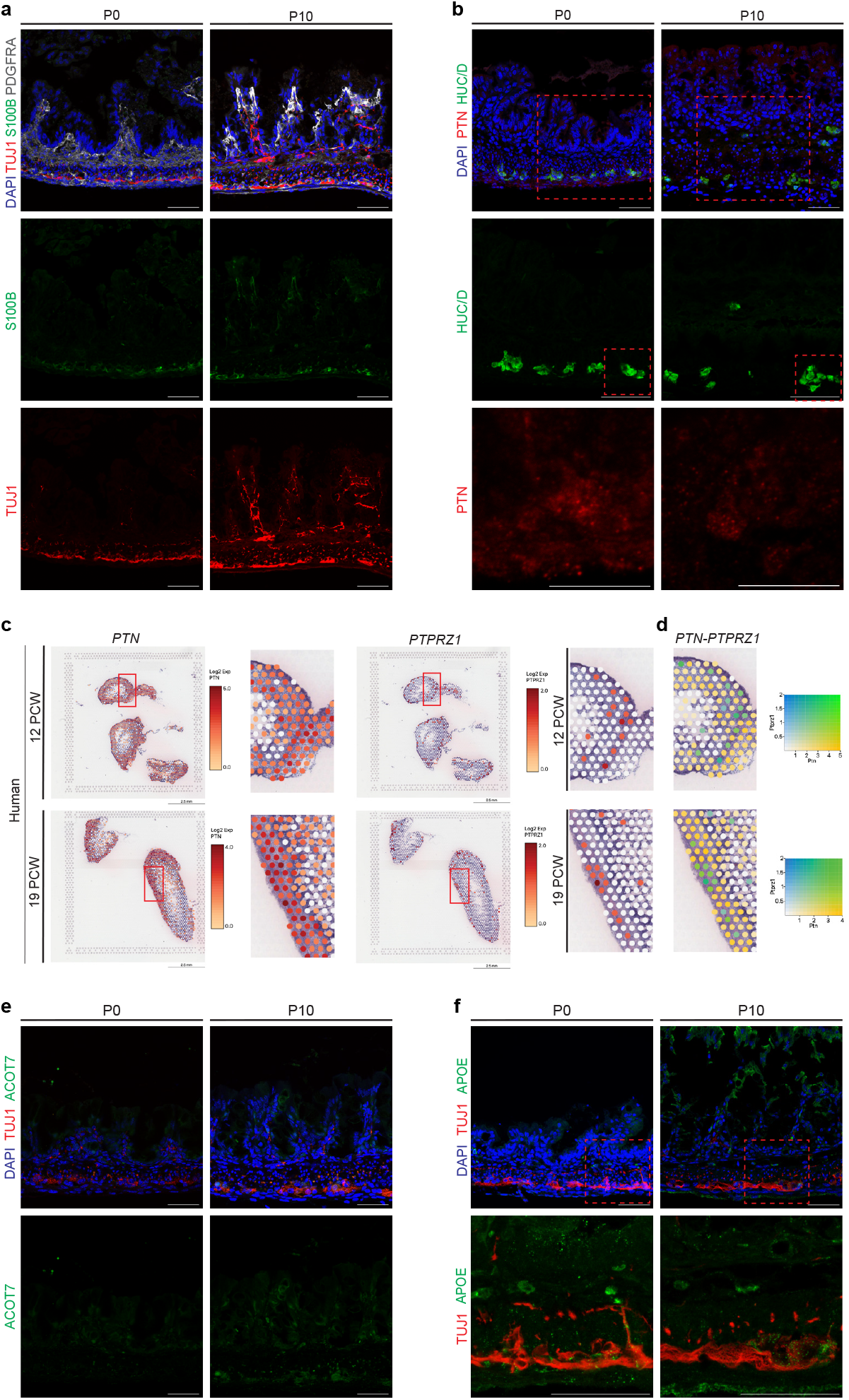
Temporal and spatial dynamics of ENS during regeneration reflect postnatal developmental patterns. **(a)** Immunohistochemistry analysis of TUJ1 (neuron), S100B (glia), and PDGFRA (mesenchymal cell) in postnatal day 0 (P0) and postnatal day 10 (P10) mouse colon. At P10, enteric glia and neurons localized to the submucosal plexus and mucosal layer. **(b)** HUC/D (neuron) and PTN histological analysis for P0 and P10 mouse colon. **(c, d)** Spatial transcriptomics analysis (GEO accession GSE158328) of 12 post-conceptional weeks (12 PCW) and 19 post-conceptional weeks (19 PCW) human data. *PTN* and *PTPRZ1* mRNA were localized in the muscle layer. *PTN-PTPRZ1* co-expression analysis indicated ligand-receptor interaction at both 12 and 19 PCW timepoints. **(e)** Histological analysis of newborn mice at P0 and P10 using ACOT7. ACOT7 expression was seen in the myenteric plexus at P10, co-expressed with TUJ1. **(f)** Histological analysis for APOE in P0 and P10 colon. APOE was detected in the myenteric plexus of the newborn mouse colon. At least 3 biological replicates (n=3) were used for tissue staining. Scale bar: 50 µm.

Similarly, spatial transcriptomics^50^ of human colonic development showed a progressive increase in *PTN* and *PTPRZ1* expression, particularly within the myenteric plexus, by postconceptional week (PCW) 19. This organization contrasted with the more diffuse expression observed at PCW 12, reflecting a developmental maturation of ENS alignment and signaling (Fig. 4c). Co-expression analysis further indicated that *PTN* and *PTPRZ1* expression became confined to the myenteric plexus in PCW 19 (Fig. 4d). Genes involved in cell adhesion and migration, including *NCAM1, L1CAM*, and *MCAM* expressions localized more prominently in the myenteric plexus during this period, paralleling the recovery phase in the colitis model (Extended Data Fig. 6a).

In light of the observed changes in lipid metabolism during murine colon regeneration, it was compelling to further investigate its role in development. ACOT7 expression, absent at P0, became prominent in the myenteric plexus at P10 (Fig. 4e). APOE expression was observed in TUJ1+ neurons within the myenteric plexus at both P0 and P10, further underscoring its role in ENS function during development (Fig. 4f). Additionally, we observed patterned expression of *PTGS2* within the myenteric plexus during human development, aligning with its role in murine tissue regeneration (Extended Data Fig. 6b). The conserved mechanisms of structural reorganization and metabolic adaptation during mammalian colon development and regeneration suggest shared regulatory mechanisms that govern ENS elongation and migration.

## Discussion

Our findings establish ERAs as a novel platform for modeling the enteric nervous system and its responses to environmental and physiological challenges. Unlike traditional organoids, these assembloids incorporate diverse cellular components, including mesenchymal cells and ENS, which self-organize into structures that closely mimic native colonic tissue. This intrinsic capacity for self-organization highlights the unique potential of assembloids to model complex multicellular interactions, positioning them as a powerful tool for understanding ENS-driven tissue dynamics. The identification of the PTN-PTPRZ1 signaling axis as a central pathway in ENS recovery underscores the importance of conserved molecular mechanisms governing neural-glial interactions. Moreover, our discovery of lipid transfer from enteric neurons to glial cells during tissue repair highlights a pioneering mechanism of metabolic support. This lipid exchange may mitigate neural stress and preserve ENS functionality, with broader implications for understanding similar processes in the peripheral nervous system^51-53^. Future studies should explore the therapeutic potential of modulating lipid metabolism to enhance neural resilience in both enteric and central nervous systems. Parallels between tissue recovery and early postnatal development reveal conserved pathways regulating cellular migration and elongation. By bridging in vivo and in vitro models, this study advances our understanding of ENS dynamics and introduces a versatile platform for translational research^54-56^. ERAs hold promise for a wide range of applications, from drug discovery to personalized medicine, particularly in conditions involving gut-brain axis dysregulation, inflammatory diseases, and neurodegeneration.

## Materials and Methods

### Mice

Animal experiments were conducted in compliance with protocols approved by the Animal Ethics Committee at Bilkent University (2019/18, 2023/17 and 2023/18). C57BL/6 and FVB/N mice, aged 8–12 weeks, of both sexes, were bred, housed, and handled at the Bilkent University Animal Facility. Animals were provided food and water ad libitum. All mice were sourced from the institution’s live mouse repository.

### DSS-Induced Colitis Animal Model

Mouse colitis was induced by administering 2.5% dextran sulfate sodium (DSS, MP Biomedicals, CA, USA, 02160110) dissolved in drinking water, provided ad libitum for five days. Mice sacrificed at the end of this period were designated as the ‘no recovery’ group. In other groups, after the five-day DSS treatment, the drinking water was replaced with regular water for two days. These mice were sacrificed at the end of the respective recovery period and classified as ‘2-day recovery’ group. Control mice received regular drinking water for 5–7 days.

### Tissue Collection and Preparation for Histological Analysis

Colon tissues were flushed with ice-cold phosphate-buffered saline (DPBS, Gibco, #14190144) to remove fecal content and divided into proximal and distal sections, each cut into 1 cm pieces. The tissues were washed twice with 1X DPBS for 5 minutes each. For paraffin embedding, the tissues were fixed in 4% paraformaldehyde (PFA, Fluka, 76240) overnight at 4°C, then washed and dehydrated sequentially with 30% ethanol/DPBS and 50% ethanol/ddH_2_O for 1 hour at room temperature. Afterward, the tissues were placed in 70% ethanol before embedding in paraffin cassettes. A Leica Automatic Benchtop Tissue Processor was used for alcohol dehydration and the waxing procedure, followed by embedding in paraffin using a Leica Paraffin Embedding Workstation. The tissues were left to solidify for 1 hour at +4°C and then stored at room temperature. For frozen embedding, the tissues were fixed in 4% PFA on ice for 1 hour, washed with DPBS, incubated overnight in 30% sucrose (Fisher Chemicals, NH, USA, #10634932) at 4°C, embedded in Tissue-Tek O.C.T. compound (Sakura Finetek, #4583), and stored at -80°C.

### Immunohistochemical Staining Protocols

For paraffin-embedded tissues, paraffin blocks were sectioned at a thickness of 5 µm and mounted onto Superfrost Plus slides (Mensel Glaser, #J1800AMNZ). Slides were placed on a stretching table at 38°C for at least 16 hours. Dewaxing was performed using two xylene (ISOLAB, #9900152501) incubations of 7.5 minutes each, followed by rehydration through a graded ethanol series (100%, 96%, 80%, 70%, and 50%), with two incubations for each concentration, lasting 4 minutes each at room temperature (RT). The sections were washed with 1X PBS (in-house recipe) three times for 5 minutes each. Antigen retrieval was performed with citrate or tris buffer (in-house recipe) under pressurized steam for 35 minutes, followed by cooling for 20 minutes and two additional washes with 1X PBS for 5 minutes each. Blocking buffer containing 10% donkey serum (Biowest, #S2170-050), 0.3% Triton X-100 (Sigma, #T-8787), and 10% 10X PBS (in-house recipe) diluted in double-distilled water was applied for 1 hour at RT. Primary antibody solution prepared in blocking buffer was applied, and the slides were incubated overnight in a humidified chamber at 4°C. Following primary antibodies were used for staining: Anti-ACOT7 antibody (Sigma, Rabbit Polyclonal, HPA025735, 1:100), anti-E.Cadherin antibody (BD Biosciences, Mouse Monoclonal, #610182, 1:50), anti-KI67 antibody (Abcam, Rabbit Monoclonal, #AB16667, 1:100), anti-LYVE1 antibody (Thermo Fisher Scientific, Rat Monoclonal, #14-0443-82, 1:200), anti-HUC/D antibody (Invitrogen, Mouse Monoclonal, #A-21271, 1:100), anti-PDGFRA antibody (R&D Systems, Goat Polyclonal, #AF1062, 1:75), anti-PHOX2B antibody (R&D Systems, Goat Polyclonal, #AF4940, 1:100), anti-SOX9 antibody (Millipore, Rabbit Polyclonal, #AB5535, 1:100), anti-TUBB3 antibody (Biolegend, Rabbit Polyclonal, #PRB-435P, 1:100), anti-GFAP antibody (Invitrogen, Rat Monoclonal, #13-0300, 1:75), anti-TUBB3 antibody (Abcam, Mouse Monoclonal, 1:200), anti-Vimentin antibody (Thermo Fisher Scientific, Rabbit Polyclonal, #PA527231, 1:100), anti-APOE antibody (Abcam, Rabbit Monoclonal, #AB183597, 1:100), anti-PTN antibody (Proteintech, Rabbit Polyclonal, #27117-1-AP, 1:100), anti-PTPRZ1 antibody (Bioss, Rabbit Polyclonal, #bs-11-327, 1:100), anti-MUC2 antibody (Abcam, Rabbit Polyclonal, #AB97386, 1:100), anti-ACTA2 antibody (Sigma, Mouse Monoclonal, #A2547, 1:100). The slides were then washed with 1X PBS three times for 5 minutes each at RT and incubated with a secondary antibody solution diluted in blocking buffer for 1 hour at RT. Following secondary antibodies were used for staining: Cy2 AffiniPure Donkey Anti-Rat IgG (H+L) (Jackson Immunoresearch Europe, Donkey Polyclonal, #712-225-153, 1:400), Cy2 AffiniPure Donkey Anti-Goat IgG (H+L) (Jackson Immunoresearch Europe, Donkey Polyclonal, #705-225-147, 1:400), Cy2 AffiniPure Donkey Anti-Mouse IgG (H+L) (Jackson Immunoresearch Europe, Donkey Polyclonal, #715-225-151, 1:400), Cy3 AffiniPure Donkey Anti-Rat IgG (H+L) (Jackson Immunoresearch Europe, Donkey Polyclonal, #712-165-153, 1:400), Cy3 AffiniPure Donkey Anti-Mouse IgG (H+L) (Jackson Immunoresearch Europe, Donkey Polyclonal, #715-165-151, 1:400), Cy3 AffiniPure Donkey Anti-Rabbit IgG H+L) (Jackson Immunoresearch Europe, Donkey Polyclonal, #711-165-152, 1:400), Cy5 AffiniPure Donkey Anti-Goat IgG (H+L) (Jackson Immunoresearch Europe, Donkey Polyclonal, #705-175-147, 1:400), Cy5 AffiniPure Donkey Anti-Rabbit IgG (H+L) (Jackson Immunoresearch Europe, Donkey Polyclonal, #711-175-152, 1:400), Cy5 AffiniPure Donkey Anti-Rat IgG (H+L) (Jackson Immunoresearch Europe, Donkey Polyclonal, #712-175-153, 1:400). Subsequent washes included two rinses with 1X PBS and two with PBST (PBS/0.03 % Triton X-100) for 5 minutes each. Nuclei were stained with DAPI (Thermo Fisher Scientific, #62248) (1:1500 in PBST) for 10 minutes at RT, followed by two washes with PBST and two with 1X PBS, each for 5 minutes. For frozen blocks, tissue sections (5–15 µm) were mounted onto Superfrost Plus slides and placed on a stretching table at 38°C for 30 minutes, then left at RT for another 30 minutes. Slides were fixed in the appropriate fixative for 10 minutes, followed by three washes with 1X PBS for 5 minutes each. Blocking buffer was applied for 1 hour at RT, and slides were incubated with the primary antibody solution overnight at 4°C. Following primary antibodies were used for staining: anti-S100B antibody (Abcam, Rabbit Monoclonal, #AB52642, 1:100), anti-PDGFRA antibody (R&D Systems, Goat Polyclonal, #AF1062, 1:75), anti-TUBB3 antibody (Biolegend, Rabbit Polyclonal, #PRB-435P, 1:100), anti-TUBB3 antibody (Abcam, Mouse Monoclonal, 1:200). Secondary antibody staining procedure and antibodies with same dilution were used as described above. Nuclei were stained with DAPI (Thermo Fisher Scientific, #62248) (1:1500 in PBST) for 10 minutes at RT, followed by two washes with PBST and two with 1XPBS, each for 5 minutes. Slides were mounted using anti-fade fluorescence mounting medium (Abcam, #AB104135), covered with a coverslip (EPREDIA, #6205119), and stored at 4°C until analysis. Images were acquired using a Leica Confocal SP8 microscope and LAS X software, utilizing 20X, 40X, and 63X objectives. Quantification of fluorescence intensity was performed with ImageJ software, and data analysis was conducted using GraphPad Prism software (9.4.0).

### Preparation of Coated Coverslips for Cell Culture

Round coverslips (18 mm) (EPREDIA, #11709875) were sterilized by submerging them in 100% ethanol for 15 minutes and then air-dried in a cell culture hood for an additional 15 minutes. Sterile forceps were used to place the coverslips into a 12-well plate (Greiner, #665180). For the lipid transfer assay, small pieces of sterilized dental wax (Integra, #1.1023 Lb) treated with 100% ethanol and UV-irradiated, were placed onto the coverslips to create a gap for the glia-neuron sandwich culture system. To prepare the coverslips, a 100 µg/mL Poly-D-Lysine (PDL) (Sigma, #P6407) solution, diluted in sterile water, was applied to cover the glass surface. For quick coating, coverslips were incubated with PDL at room temperature for at least 1 hour. For long-term coating, the PDL solution was left on the coverslips overnight at +4°C in a securely sealed cell culture plate covered with parafilm. After incubation, the PDL solution was carefully removed without disturbing the glass surface, and the coverslips were allowed to air-dry completely in the culture hood. Once dry, a 10 µg/mL laminin (Sigma, L2020) solution diluted in sterile PBS was applied to the PDL-coated coverslips, covering only the glass surface. The laminin-coated coverslips were incubated at 37°C for 2 hours. After incubation, the laminin solution was removed without scratching the surface, and the wells were gently washed three times with 1X PBS. Respective media were added to each well, and the plates were stored at 37°C until cell seeding.

### ENS Cell Isolation for Lipid Transfer

The colon was dissected into ice-cold HBSS (Gibco, #24020-117), and the longitudinal muscle and myenteric plexus (LMMP) layers were peeled off. The LMMP was immediately placed into ice-cold HBSS. From this point, glia and neuron isolation protocols proceeded separately. For glia isolation, the LMMP was cut into tiny pieces in 5 mL Corning Cell Recovery Solution (Corning, #354253) and incubated on a rotating platform for 30 minutes at +4°C. The solution containing LMMP pieces was further dissociated by triturating 20 times with a 5 mL serological pipette coated with 0.1% BSA/PBS (Sigma, #A7030). To recover LMMP pieces, the solution was filtered through a 100 µm strainer (Greiner, #542000) and washed with 2 mL of 1X DPBS (Gibco, #14190144). The supernatant was discarded, and the tissue pieces retained on the strainer were transferred with clean forceps into 1.5 mL of digestion medium consisting of HBSS (Gibco, #24020-117) with 10 mM HEPES (Gibco, 15630056), 50 µg/mL Liberase TM (Roche, #5401119001), and 100 U/mL DNase 1 (Sigma, #DN25) in a 2 mL Eppendorf tube. This mixture was incubated on a rotating platform for 30 minutes at 37°C. Enzymatic activity was stopped by adding complete media, and the solution was immediately filtered through a 70 µm strainer. The supernatant was transferred to a 15 mL tube, brought to a volume of 5 mL with complete media, and centrifuged at +4°C at 600×g for 5 minutes. The supernatant was carefully discarded, and the pellet was washed with 1X DPBS, followed by centrifugation at +4°C at 600×g for 5 minutes. The cells were counted, and 150,000 cells/well were seeded on dental wax-attached coverslips in a 12-well plate. For neuron isolation, the LMMP was cut into tiny pieces in digestion solution consisting of HBSS (Gibco, #24020-117) with 10 mM HEPES (Gibco, 15630056), 100 µg/mL Liberase TM (Roche, #5401119001), and 100 U/mL DNase 1 (Sigma, #DN25) in a 2 mL Eppendorf tube and incubated on a rotating platform for 30 minutes at 37°C. The tissue pieces were triturated 10 times and filtered through a 100 µm strainer. Enzymatic activity was stopped by adding complete media to the supernatant, which was then centrifuged at 600×g for 5 minutes at 4°C. The pellet was washed once, resuspended in complete media, and counted. A total of 150,000 cells/well were seeded on coverslips in a 12-well plate. Cells were maintained in complete media consisting of DMEM/F12 with L-Glu (Capricorn Scientific, #DMEM-12-A), 15 mM HEPES (Gibco, 15630056), 100 U/mL Penicillin/Streptomycin (Gibco, #15140148), and 10% FBS (Gibco, #16140071) for 2 days. At day 3, glia differentiation was induced using a new medium consisting of DMEM/F12 with L-Glu (Capricorn Scientific, #DMEM-12-A), 15 mM HEPES (Gibco, 15630056), 100 U/mL Penicillin/Streptomycin (Gibco, #15140148), 0.2% N2 (Gibco, #17502048), and 10 µg/mL GDNF (PeproTech, #450-44). At day 3, neurons were differentiated using NeuroBasal Plus medium (NB+) (Gibco, #A35829-01), 2% B27 Plus, and 0.5 mM GlutaMAX I Supplement (Gibco, #35050038). Half of the media was renewed every 2 days. Glia and neuron cells were used for the lipid transfer protocol 5 days after the initial seeding.

### Lipid Transfer Using ENS Cells

On day 5 of culture, glial cells in the treatment group were exposed to 10 ng/mL IL-1β (Peprotech, #211-11B), TNF-α (Peprotech, #315-01A), and IFN-γ (Peprotech, #315-05) for 48 hours by changing half of their media. Neurons were treated with 2.5 mM C12 (Invitrogen, #D3835) for lipid labeling 18 hours before lipid transfer by replacing half the medium (1 mL). On day 7, after 18 hours of C12 labeling, neurons were washed twice with pre-warmed DPBS (Gibco, #14190144) and incubated in fresh NeuroBasal Plus (NB+) (Gibco, #A35829-01) medium for 1 hour at 37°C. Both neuron and glial wells were washed twice with pre-warmed DPBS. Pre-warmed HBSS-calcium chloride medium (Ca/Mg-free HBSS without phenol red (Capricorn Scientific, #HBSS-2A), 20 mM HEPES (Gibco, #15630056), 2 M calcium chloride (Sigma, #C3306) was added to the glial wells. The neuronal coverslip was inverted, placed on top of the glial coverslip, stabilized on dental wax to form a sandwich culture, and incubated at 37°C for 4 hours. After incubation, the neuronal coverslip was carefully removed and returned to its original well. Lipid transfer was imaged by staining nuclei with DAPI (Thermo Fisher Scientific, #62248, 1:3000) to preserve C12 signal intensity. Coverslips were washed twice with DPBS (5 minutes each), fixed with 3% PFA for 15 minutes at room temperature, washed again with PBS, and stained with DAPI (Thermo Fisher Scientific, #62248, 1:3000). Coverslips were mounted with anti-fade fluorescence mounting medium (Abcam, #AB104135) and imaged immediately using Leica confocal microscope. To ensure neuronal and cell-type purity, coverslips were stained following the same fixation steps. For the primary antibody incubation, a primary antibody solution prepared in blocking buffer (2% BSA (Sigma, #A7030) and 0.2% Triton X-100 (Sigma, #T-8787) in PBS) was applied to a parafilm-covered slide. Following primary antibodies were used for staining: Anti-S100B antibody (Abcam, Rabbit Monoclonal, #AB52632, 1:100), anti-GFAP antibody (Invitrogen, Rat Monoclonal, #13-0300, 1:75), anti-TUBB3 antibody (Biolegend, Rabbit Polyclonal, #PRB-435P, 1:100), anti-TUBB3 antibody (Abcam, Mouse Monoclonal, 1:200). Coverslips containing cells were inverted onto the solution and incubated for 1 hour. After incubation, coverslips were washed three times for 15 minutes each in wash buffer (0.1% Triton X-100 in PBS) on a shaker. The same procedure was repeated for the secondary antibody incubation with same antibodies mentioned. To remove unbound antibodies, coverslips were washed three times for 15 minutes each in wash buffer. Nuclei were stained with DAPI (Thermo Fisher Scientific, #62248, 1:3000) for 15 minutes. Coverslips were then washed three times for 15 minutes each in wash buffer, mounted using an anti-fade fluorescence mounting medium, and imaged using a Leica confocal microscope.

### ENS Cell Isolation for Assembloid Culture

The colon tissues were dissected into ice-cold 1X DPBS (Gibco, #14190144). The longitudinal muscle and myenteric plexus (LMMP) layers were peeled. The LMMP was immediately transferred to ice-cold DPBS and cut into 2-4 mm^2^ pieces in a 2 mL Eppendorf tube containing a digestion solution of 50 U/mL penicillin-streptomycin (Gibco, #15140148) and 0.1 mg/mL Liberase TM (Roche, #5401119001) in HBSS without Ca/Mg (Gibco, #14170088). The samples were incubated on a rotator at 37°C for 30 minutes. After incubation, the Eppendorf tube was gently shaken, and the tissue was triturated 15 times. The solution was briefly centrifuged to separate the LMMP pieces from the supernatant. The supernatant was transferred to a fresh 1.5 mL Eppendorf tube, and FBS (Gibco, #16140071) was added to halt enzymatic activity. The LMMP pieces remaining in the pellet were subjected to further digestion with 0.05% Trypsin-EDTA solution (Gibco, #25300-054) at 37°C for 10 minutes on a rotator. Following digestion, the solution was triturated 15 times. Both the Trypsin-EDTA-treated supernatant and the earlier collected supernatant were filtered through a 70 µm strainer into a Falcon tube and centrifuged at 150×g for 5 minutes in Advanced DMEM/F12 (Gibco, #12634028). The resulting cell pellet was resuspended, and cells were counted. A total of 150,000 cells were seeded per well in a 12-well plate (Greiner, #665180). Cells were maintained in glial cell media consisting of Advanced DMEM/F12 supplemented with 100 U/mL penicillin-streptomycin (Gibco, #15140148), 10% FBS, 20 µg/mL gentamicin (Sigma, #G1397), 1X GlutaMAX I Supplement (Gibco, #35050038), and 10 µg/mL GDNF (Peprotech, #450-44) for 2– 3 days. Subsequently, cells were differentiated into neurons using NeuroBasal Plus medium supplemented with 2% B27 Plus (Gibco, #A35829-01) and 0.5 mM GlutaMAX I Supplement. Half of the medium was replaced every 2 days. ENS cells were used for the assembloid protocol 7–8 days after initial seeding.

### Colon Crypt Isolation for Assembloid Culture

The colon was flushed with ice-cold 1X DPBS (Gibco, #14190144) and opened longitudinally. The tissue washed three times with 1X DPBS and then cut into 1 cm pieces. They were transferred into a solution of PBS containing 10 mM EDTA (Invitrogen, #15575-038) and 0.5 mM DTT (Sigma, #D9779). Tissue dissociation was carried out in this EDTA/DTT solution on a rotator at 37°C for 20 minutes. Following incubation, the solution was replaced with ice-cold 1X DPBS, and the tissue pieces were shaken vigorously for 30 seconds to release crypts. The supernatant containing crypts was collected and centrifuged at 400×g for 5 minutes at 4°C. The remaining tissue pieces were reserved for mesenchymal cell isolation. Crypts were counted and seeded at a density of 150 crypts per dome into 10 µL of matrigel (Corning, #354255). Three matrigel domes were plated per well in a 24-well plate (Greiner, #662160). The matrigel domes were polymerized at 37°C for 10 minutes and then overlaid with organoid growth medium. The organoid growth medium consisted of Advanced DMEM/F12 (Gibco, #12634028) supplemented with 10 mM HEPES (Gibco, #15630056), 2 mM GlutaMAX I Supplement (Gibco, #35050038), 100 U/mL penicillin-streptomycin (Gibco, #15140148), 1.25 mM N-acetyl-L-cysteine (Sigma, #A2750), 1X B-27 (Gibco, #17504044), 1X N2 (Gibco, #17502048), 50 ng/mL mouse EGF (Thermo Fisher Scientific, #PMG8041), 100 ng/mL human noggin (Peprotech, #120-10C), 1 µg/mL mouse R-spondin (Peprotech, #315-32), 0.328 nM surrogate Wnt (J-Protein Express BV, #N001), 10 µM rock inhibitor Y-27632 (Sigma, #Y0503), 0.5 µM A-8301 (Tocris, #2939), and 3 µM CHIR-99021 (Biogems, #2520691). The medium was replaced every 2 days. Organoids were used in experiments after a single passage.

### Mesenchymal Cell Isolation for Assembloid Culture

The remaining tissue fragments from the crypt isolation protocol were used for mesenchymal cell isolation. The tissue fragments were washed multiple times with ice-cold DPBS (Gibco, #14190144) by vigorously shaking for 30 seconds each time until the supernatant was clear, ensuring the removal of any remaining epithelial cells. The fragments were then minced into small pieces and incubated in digestion medium containing Ca/Mg-free HBSS (Gibco, #14170088), 100 µg/mL Liberase TM (Roche, 5401119001), and 400 U/mL DNase I (Sigma, #DN25). The digestion was performed on a rotator at 37°C for 20 minutes. After digestion, the fragments were triturated to release cells, and the supernatant was collected. To stop enzymatic activity, 10% FBS (Gibco, #16140071) was added to the supernatant. This process was repeated with fresh digestion medium; the fragments were triturated again, and the supernatant was collected. The combined supernatants containing single cells were filtered through a 70 µm strainer (Corning, #352350) and centrifuged at 400×g for 5 minutes at 4°C. The resulting cell pellet was resuspended in stromal growth medium, composed of Advanced DMEM/F12 (Gibco, #12634028) supplemented with 10% FBS, 100 U/mL penicillin-streptomycin (Gibco, #15140148), and 10 µM Y-27632 (Sigma, #Y0503). The cells were seeded into a 6-well plate (Greiner, #657160), and the medium was changed every 2–3 days.

### Preparation and Culturing of Assembloids

Three-to four-day-old organoids were collected from Matrigel using Cell Recovery Solution (Corning, #354253) and incubated for 30 minutes to depolymerize the matrigel (Corning, #356231) completely. Organoids were washed once with ice-cold DPBS (Gibco, #14190144), centrifuged at 100×g for 1 minute, and reconstituted in matrigel at a concentration of 20 organoids per 0.5 µL. Mesenchymal cells cultured in 6-well plates were washed once with Advanced DMEM/F12 (Gibco, #12634028) and detached using 1 mL of 0.05% Trypsin-EDTA solution (Thermo Fisher Scientific, #25300-054) for 15 minutes at 37°C. Upon detachment, cells were triturated to ensure complete dissociation. The cells were collected, washed once with Advanced DMEM/F12, and centrifuged at 400×g for 5 minutes at 4°C. The pellet was resuspended in matrigel at a concentration of 1.5×10^5^ cells per 1.75 µL of matrigel. ENS cells were washed once with Advanced DMEM/F12 and detached from coverslips using 500 µL of 0.05% Trypsin-EDTA solution for 10–15 minutes at 37°C. Detached cells underwent five rounds of trituration for complete dissociation. Cells were collected, washed with Advanced DMEM/F12, and centrifuged at 400×g for 5 minutes at 4°C. The pellet was resuspended in matrigel at a concentration of 1×10^5^ cells per 1.75 µL of matrigel. A sterile parafilm sheet was placed in a 100 mm petri dish, and DPBS droplets were added to maintain humidity. ENS cells and mesenchymal cells were mixed at a 1:1 ratio, and 2 µL of the mixture was placed on the parafilm using a P2 pipette. Separately, 0.5 µL of organoids were mixed with 1.5 µL of the ENS/mesenchymal cell mixture, forming a 4 µL assembloid. The assembloids were solidified at 37°C for 30 minutes. Using fine forceps, the assembloids were transferred carefully from the parafilm to pre-warmed assembloid growth medium in a 12-well plate. Assembloids were cultured on an orbital shaker at 170 rpm in a cell culture incubator. The assembloid growth medium consisted of Advanced DMEM/F12 supplemented with 10 mM HEPES (Gibco, #15630056), 2 mM GlutaMAX I Supplement (Gibco, #35050038), 100 U/mL penicillin-streptomycin (Gibco, #15140148), 1.25 mM N-acetyl-L-cysteine (Sigma, #A2750), 1× B-27 (Gibco, #17504044), 1× N2 (Gibco, #17502048), 50 ng/mL mouse EGF (Thermo Fisher Scientific, #PMG8041), and 0.5 µM A-8301 (Tocris, #2939). Media changes were performed every 2 days. Assembloids were cultured for 8 days. For fixation, assembloids were washed once with 1× DPBS and fixed in 4% PFA (Fluka, #76241) overnight at 4°C. The following day, assembloids were washed three times with 1× PBS (in-house recipe) for 5 minutes each. To prevent adhesion to pipette tips, assembloids were incubated in 0.1% BSA/PBS (Sigma, #A7030) for 30 minutes at room temperature, followed by a final wash with 1× PBS. Assembloids were transferred to a plastic mold using a cut pipette tip, excess liquid was removed, and the assembloids were embedded in 1% agarose (Biomax, #HS-8000). For paraffin embedding, the agarose mold containing assembloids was removed from the plastic mold, and manual tissue paraffin embedding protocols were followed. For frozen sections, assembloids in the plastic mold were embedded in Tissue-Tek O.C.T. (Sakura Finetek, #4583) and stored at -20°C. Tissue sections were prepared for further analysis, and immunohistochemistry protocols were applied as appropriate.

### Single-Cell RNA Sequencing Analysis of Mouse Sample

Pre-existing single-cell RNA sequencing data from mice were analyzed using Cell Ranger version 8.0.1. A single sample, M4_S15, was selected for further analysis. The 10x Genomics BAM file for this sample was realigned to the mouse reference genome (GRCm38) using the Cell Ranger count pipeline with default parameters. The resulting .cloupe file was processed and analyzed in the Loupe Browser version 8.0.0 for data exploration and visualization. Dimensionality reduction and k-means clustering were performed within the Loupe Browser to identify distinct cell populations. Each cluster was annotated based on the expression profiles of specific marker genes. The marker genes used for annotation included the following:

Marker genes used for annotation included goblet/secretory cells (*Muc2, Villin, Fcgbp, Clca1, Atch1, Kit, Agr2, Spdef*), stem/transit-amplifying (TA) cells 1 (*Mki67, Top2a, Cenpf*), stem/TA cells 2 (*Lgr5, Clca3b*), colonocytes (*Krt20, Muc3, Slc6a14*), fibroblasts (*Pdgfra, Col1a1, Col1a2, Acta2, Rgs5*), T cells (*Trac, Trbc2, Ccl5, Klbr1, Runx3, Cd8a*), blood endothelial cells (BECs) (*Kdr, Plvap*), enteroendocrine cells (EECs) (*Cck, Chgb, Gfra3, Vwa5b2, Neurod1*), glial cells (*Sox10, GFAP*), and neurons (*Tubb3, Elavl3, Elavl4, Phox2b, Uchl1, Ret*). T-distributed stochastic neighbor embedding (t-SNE) was applied to visualize the cell clusters in two-dimensional space. Violin plots were generated in the Loupe Browser to depict the distribution of gene expression levels for marker genes across clusters. This workflow facilitated detailed annotation and characterization of the cellular composition within the selected sample.

### Spatial Transcriptomics of Inflammatory and Developmental Samples

Mouse spatial transcriptomic data from Day 0 and Day 14 and human intestinal development data from 12 post-conceptional weeks (PCW) and 19 PCW were analyzed using Space Ranger version 3.0.1 from 10x Genomics. The FASTQ files were aligned to the mouse reference genome (GRCm38) and human reference genome (GRCh37/hg19), respectively, using the Space Ranger pipeline with default parameters. The Space Ranger count pipeline was employed for analysis, with no modifications to the default settings. The resulting .cloupe files were used for visualization and downstream analysis in the Loupe Browser (10x Genomics). Tissue images were aligned with spatial transcriptomics spots using the internal alignment functionality of the Loupe Browser. For all features, scale values were transformed using a log2 scale within the Loupe Browser. In the mouse spatial transcriptomics data, specific regions were categorized into two groups based on inflammation status: “inflammatory and hyperplasia” and “mild inflammation”. Insets from the spatial transcriptomics analysis highlighted the inflammatory regions within the tissue, enabling detailed characterization of the inflammatory states in the samples.

### RNA Isolation and Quantitative RT-PCR Analysis

Total RNA was extracted from the cells isolated from muscularis externa of C57BL/6 mice at three time points: control, no recovery, and 2-day recovery. Cell lysis and RNA extraction were performed following the manufacturer’s instructions for the Qiagen RNeasy Mini Kit (Qiagen, #74104). Complementary DNA (cDNA) synthesis was conducted using the PrimeScript™ RT Master Mix (Perfect Real Time) (Takara, #RR036A), according to the manufacturer’s protocol. Quantitative real-time PCR (qRT-PCR) was performed to assess gene expression using iTaq Universal SYBR Green SMX500 (BioRad, #1725121) on the CFX96 Touch Real-Time PCR Detection System. Each reaction utilized 20 ng of cDNA as the template. β-actin was used as the internal control, and three technical replicates were included for each experimental condition. Relative gene expression changes were calculated using the 2^-ΔCt method, enabling quantification of expression differences across the experimental time points.

### Data Analysis and Statistical Methods

Quantitative data analysis was performed using GraphPad Prism software (version 9.4.0). Fluorescence intensity of protein expression was quantified using Fiji/ImageJ. For the C12 signal, quantification involved background signal subtraction for each channel, with a rolling ball radius of 20 pixels for C12 and 80 pixels for DAPI. The integrated density of the C12 signal was normalized to the integrated density of the DAPI signal. Statistical analyses included t-tests and Mann-Whitney tests for comparisons between two groups. For image-based statistical comparisons, group differences were analyzed using one-way analysis of variance (ANOVA) followed by the Kruskal-Wallis test. Data are presented as mean ± standard error of the mean (SEM). For qRT-PCR analyses, data are presented as mean ± standard deviation (SD) and analyzed using ordinary one-way ANOVA with Tukey’s multiple comparisons test. Significance thresholds were set as follows: *p < 0.05, **p< 0.01, **** p < 0.0001. Non-significant (ns) values were indicated in the corresponding graphs.

## Data Accessibility

The mouse single-cell RNA sequencing dataset utilized in this study is publicly available in the Broad Institute Single Cell Portal under the accession code SCP1038. The spatial transcriptomics datasets analyzed in this study are accessible via the following accession codes: GSE169749 (mouse DSS and recovery data) and GSE158328 (human 12PCW and 19PCW data).

## Acknowledgements

We thank M. Demirdizen, D. Esen, and all members of the BD laboratory for their valuable discussions and insights. We acknowledge the technical support provided by the staff at the Animal Facility of Bilkent University and the laboratory of M. Cevher. We also thank the members of the Basler laboratory at the University of Zurich for their advice and constructive comments during this study. This work was supported by TUBITAK (grant number 219Z163) and the EMBO Installation Grant (grant number 4442) provided to B.D.

## Author Contributions

I.S. performed and analysed most of the experiments. I.S., B.U., and N.G. conducted the mouse experiments. I.S., B.U., N.S., and E.G., carried out the 3D organoid, assembloid, and primary cell culture experiments. B.U. and M.O.O. conducted RNA isolation and qRT-PCR analyses. D.A.G. assisted with image analysis. T.V. supervised the 3D organoid and mesenchymal cell culture experiments. F.A.K., M.G., and P.K. contributed to the histology experiments. B.D. conceived the study, supervised the project, secured funding, and wrote the manuscript.

## Competing interests

The authors declare no competing interests.

**Extended Data Figure 1.**
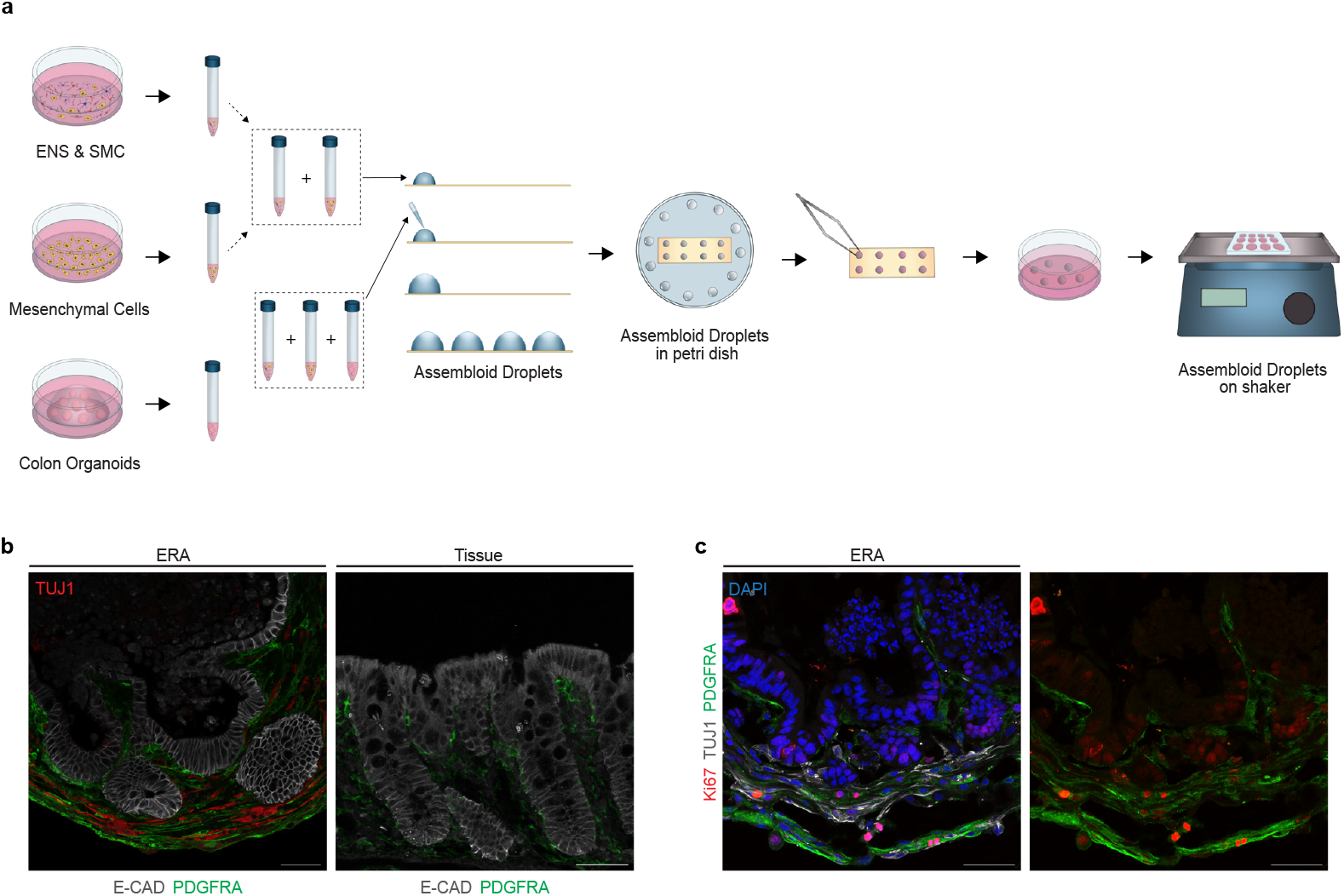
Generation of ENS-rich assembloids (ERAs) that recapitulate colon tissue architecture. **(a)** Experimental scheme for the generation of ERAs. Mesenchymal cells, ENS cells, and colon organoids were isolated and plated separately, then combined in a Matrigel droplet to form ERAs. **(b)** Histological comparison between ERAs and native colon tissue. Immunostaining of E-CAD (epithelial cells), TUJ1 (neurons), and PDGFRA (mesenchymal cells) reveals similar tissue architecture. PDGFRA+ cells are localized around crypt-like structures in the assembloids, mirroring native colon architecture. **(c)** Histological analysis of ERAs with cell-specific markers. Both crypt base epithelial cells and PDGFRA+ mesenchymal cells exhibit active proliferation, as indicated by Ki67 staining. TUJ1 stains the neurons present in the assembloids. Representative data from at least three biological replicates (n=3), with three assembloids per experimental group. Scale bar: 50 µm.

**Extended Data Figure 2.**
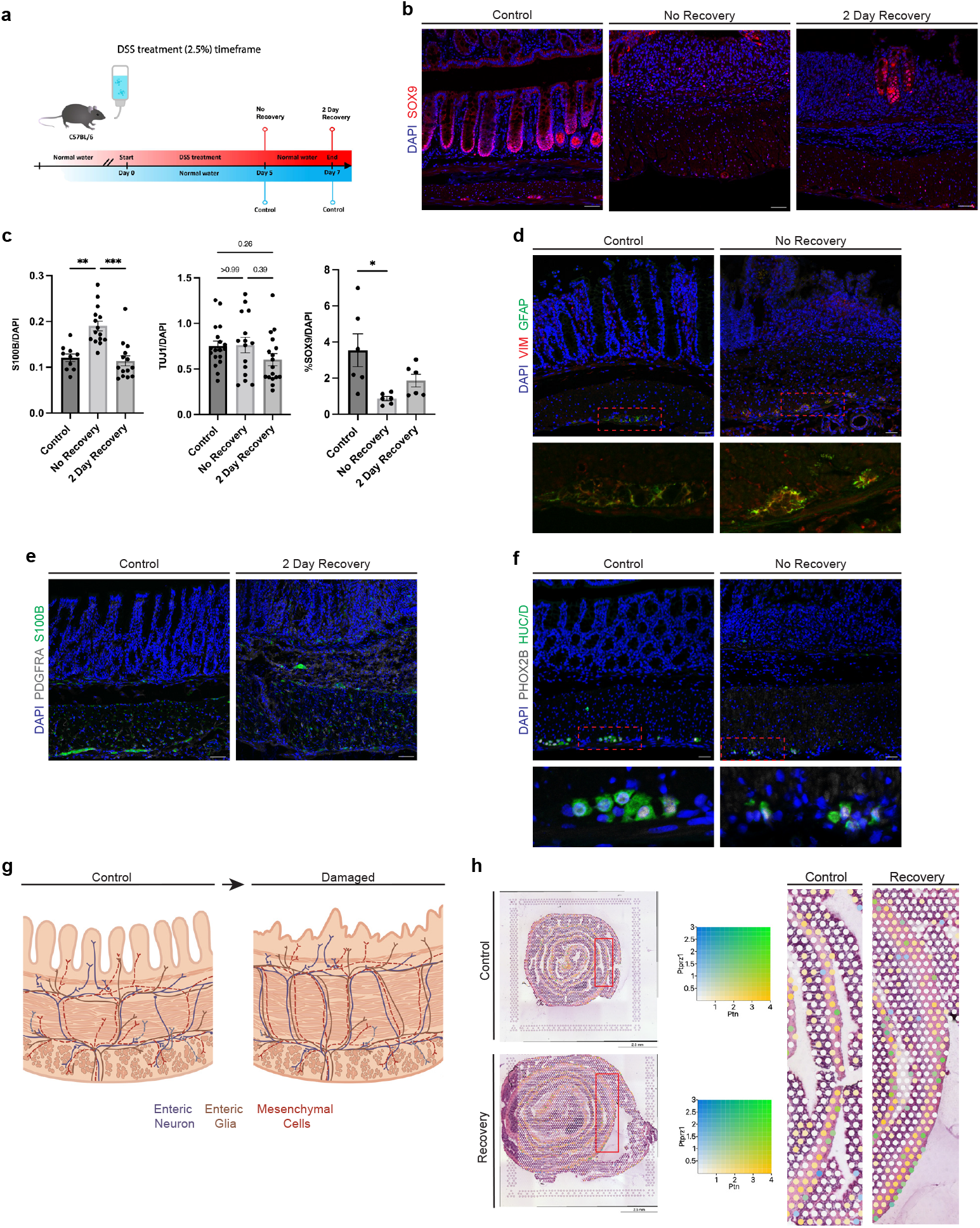
Spatial dynamics of ENS and mesenchymal cells during damage and recovery. **(a)** Schematic of the colitis model. **(b)** Immunohistochemistry analysis of the progenitor marker SOX9 in colon tissue from control, no recovery, and two-day recovery groups. **(c)** Quantification of S100B (glial marker), TUJ1 (neuronal marker), and SOX9 expression in colon muscle area. Graphs show mean ± SEM. *p < 0.05, **p < 0.01, ****p < 0.0001. **(d)** GFAP (glia) expression overlapped with VIMENTIN (VIM) in the myenteric plexus. **(e)** Staining for S100B and PDGFRA demonstrated that expression of these markers returned to baseline levels after 2 days of recovery. **(f)** Neuronal markers HUC/D and PHOX2B were used to characterize neuron abundance in the myenteric plexus. **(g)** Schematic representation of increased coupling between ENS cells and mesenchymal cells upon damage. **(h)** Co-expression analysis of *Ptn* and *Ptprz1* in control and recovery tissues using spatial transcriptomics (GEO accession GSE169749). Increased co-localization of *Ptn* and *Ptprz1* was observed in the recovery site (inset). At least 3 biological replicates (n=3) were used for tissue staining and image analysis. Scale bar: 50 µm.

**Extended Data Figure 3.**
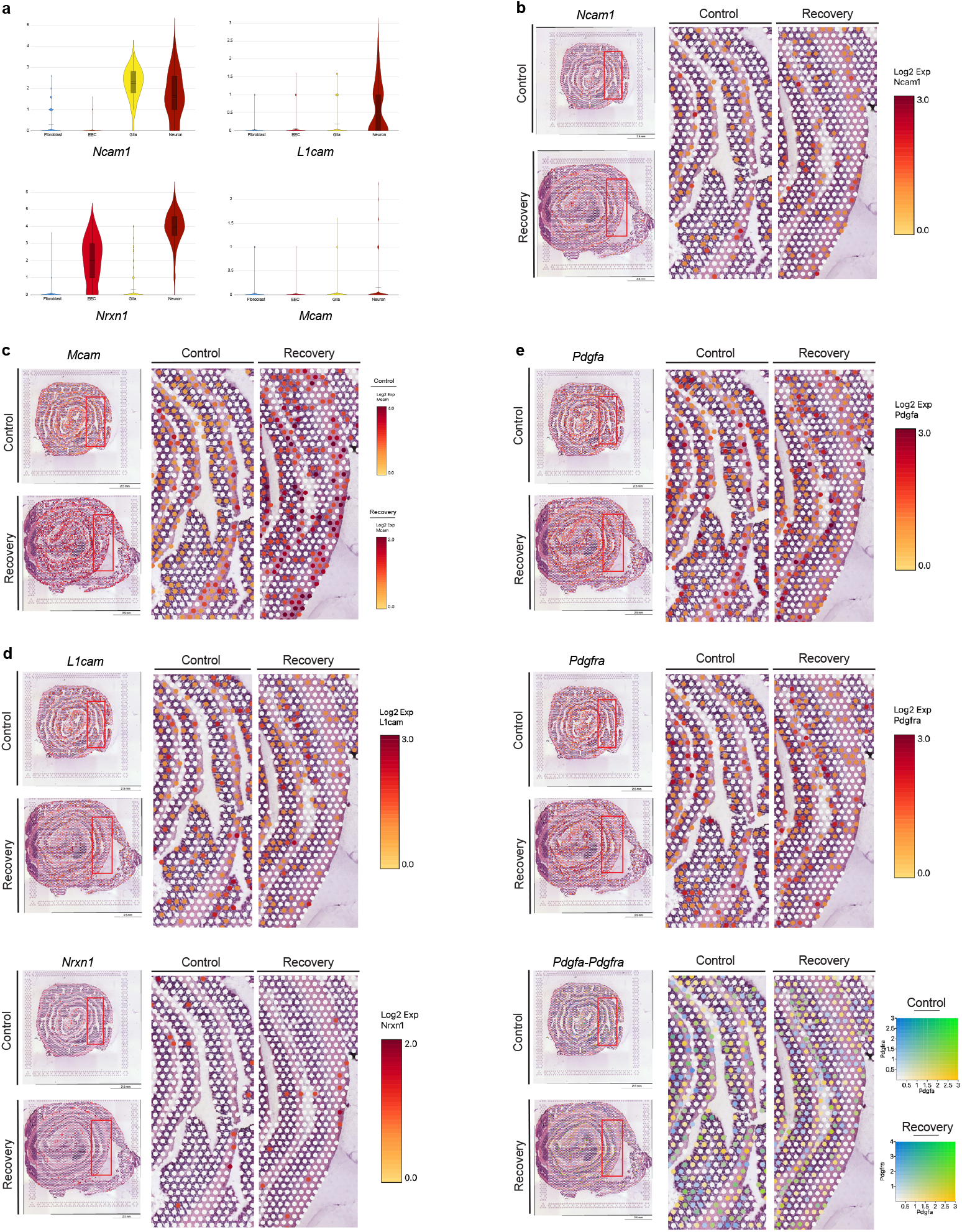
Expression profiles of healthy and regenerated colon reveals key genes mediating ENS inclination. **(a)** Violin plots show expression of *Ncam1, L1cam, Nrxn1* and *Mcam* in scRNA-seq data (Broad Institute Single Cell Portal Accession Number SCP1038). **(b-e)** Spatial transcriptomics analysis (GEO accession number GSE169749) of genes *Ncam1, L1cam, Nrxn1, Mcam, Pdgfra* and *Pdgfa*.

**Extended Data Figure 4.**
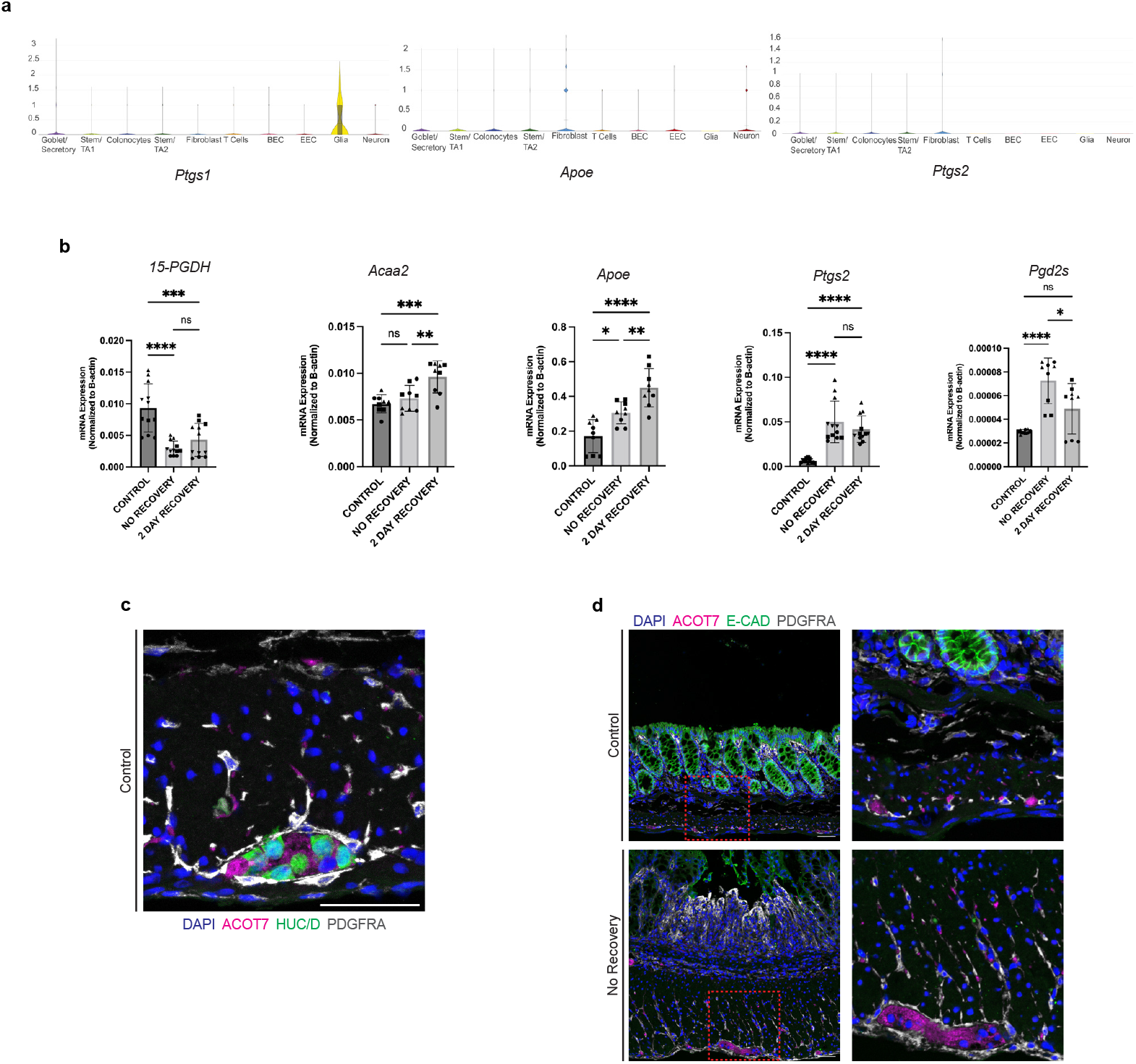
Alterations in lipid metabolism genes in response to inflammation and regeneration. **(a)** Violin plots from the analysis of single-cell RNA sequencing data (Broad Institute Single Cell Portal Accession Number SCP1038) of healthy mouse colon show the expression patterns of *Ptgs1, Ptgs2*, and *Apoe*. **(b)** qRT-PCR analysis of fat metabolism related genes. Change in gene expression levels of *15-PGDH, Acaa2, Apoe, Ptgs2, Pgd2s*; each point represents technical replicate of three biological replicates. Data presented as mean ± SD and analyzed with ordinary one way ANOVA with Tukey’s multiple comparisons test *p < 0.05, **p < 0.01, ****p < 0.0001. **(c)** Immunohistochemistry analysis of ACOT7, PDGFRA (mesenchymal cell) and HUC/D (neuron) in healthy colon. **(d)** Histological analysis of ACOT7, PDGFRA and E-CAD (epithelial cell) in healthy and damaged colon. At least 3 biological replicates (n=3) were used for tissue staining and qRT-PCR analysis. Scale bar: 50 µm.

**Extended Data Figure 5.**
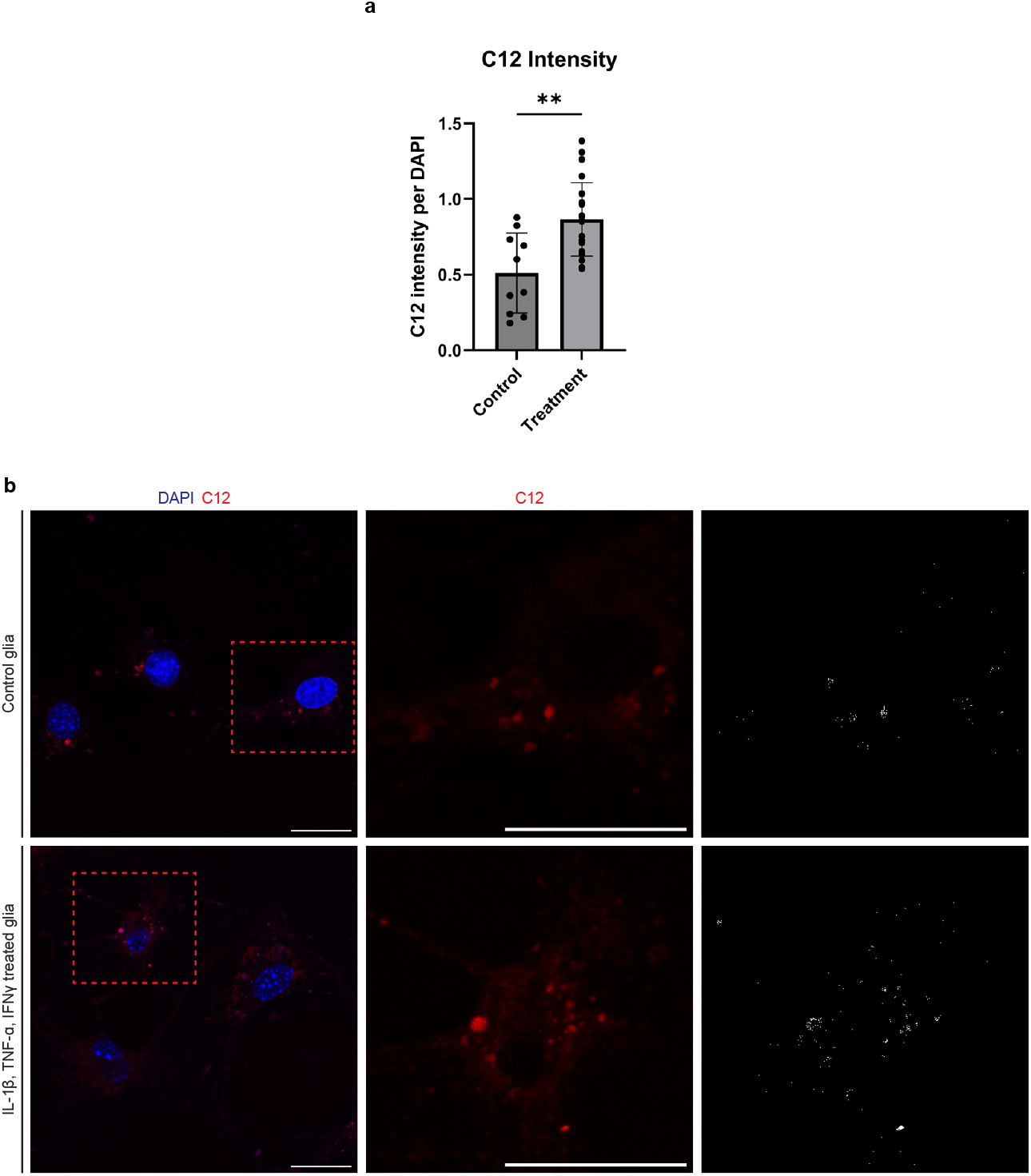
Lipid transfer experiment from enteric neuron to glia demonstrates enteric glia as a protective hub for enteric neurons. **(a)** Quantification of signal intensity of C12. Graphs are reported as mean ±SD. Each point represent quantification from one image. *p < 0.05, **p < 0.01, ****p < 0.0001. **(b)** Different set of sandwich coculture experiment with non-treated (control) glia and IL-1β, TNF-α, IFN-γ treated glia revealed C12 lipid transfer from neuron to glia and lipid transfer increase upon treatment. At least 3 biological replicates (n=3) were used for cell staining. Scale bar: 50 µm.

**Extended Data Figure 6.**
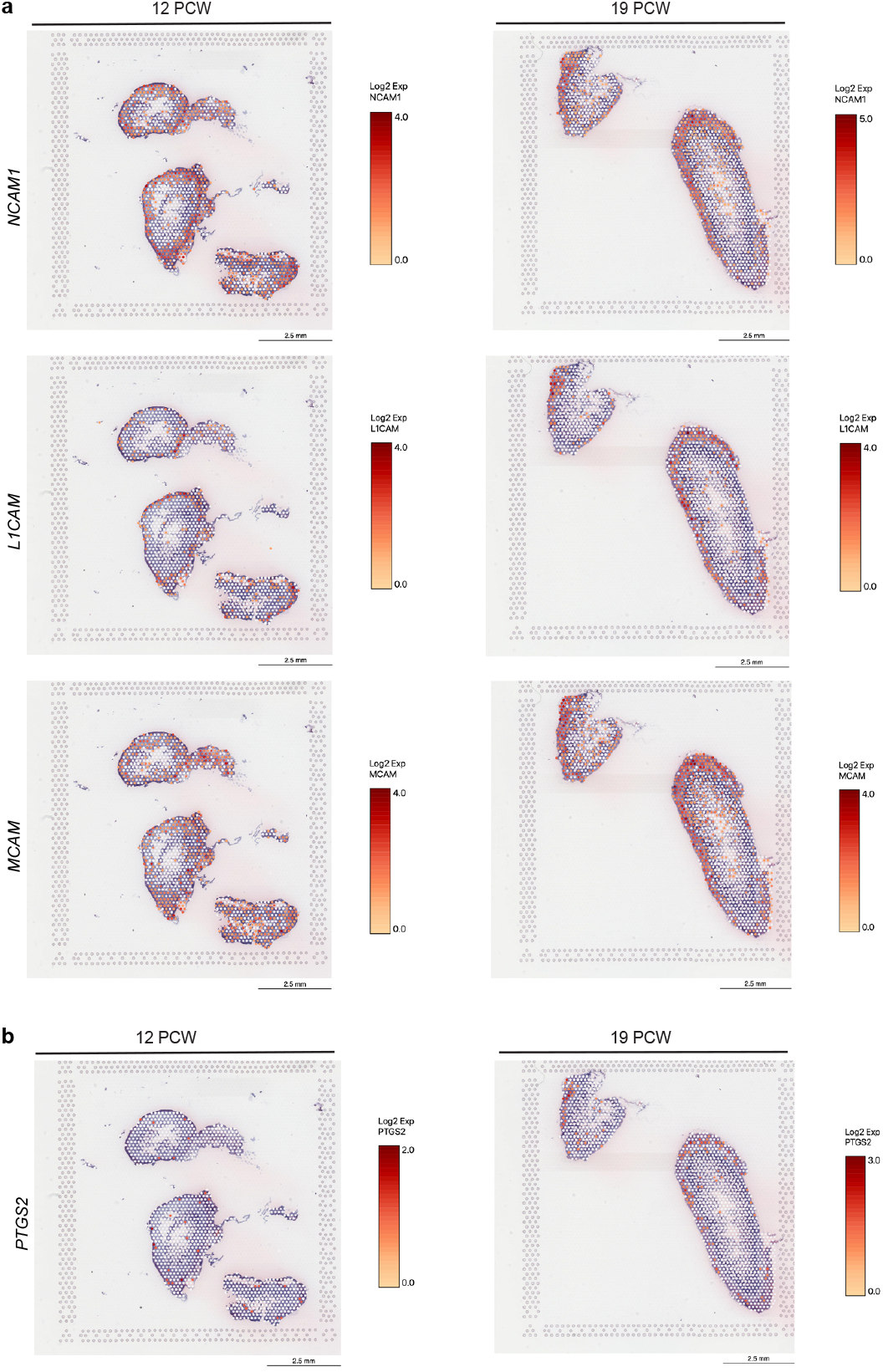
Spatial transcriptomics analysis of human 12-19 PCW reveals selective localization of genes involved in ENS development. **(a)** Spatial transcriptomics analysis (GEO accession GSE158328) of human samples at 12 PCW and 19 PCW identified the localization patterns of genes *NCAM1, L1CAM*, and *MCAM*. **(b)** Further analysis of the same dataset (GEO accession GSE158328) showed the spatial distribution of *PTGS2* at 12 PCW and 19 PCW.

## Notes

### Competing Interest Statement

The authors have declared no competing interest.

